# Comparative distribution of antisense-RNA regulated toxin-antitoxin systems

**DOI:** 10.1101/060863

**Authors:** Dorien S Coray, Nicole Wheeler, Jack A Heinemann, Paul P Gardner

## Abstract

Toxin-antitoxin (TA) systems are gene modules that appear to be widely horizontally mobile. It has been proposed that type I TA systems, with an antisense RNA-antitoxin, are less mobile than other TAs but no direct comparisons have been made. We searched for type I, II and III toxin families on chromosomes, plasmids and phages across bacterial phyla.The distribution of type I TA systems were more narrow than most type II and III system families, though this was less true of more recently discovered families.We discuss how the function and phenotypes of type I TA systems as well as biases in our databases and discovery of these modules may account for differences in their distribution.

## Introduction

Type I toxin-antitoxin (TA) systems are two-gene modules comprised of a toxic protein and an antisense RNA antitoxin. They are one of five types of TA systems, grouped according to the mechanism of the antitoxin and whether it is a protein or an RNA molecule. Of the five types of TA systems, the first and best-described are type I and type II, which encode a protein antitoxin that directly binds the toxin protein. The more recently described type III TA systems have an RNA antitoxin that directly binds the protein ^1^. The type I TA RNA antitoxins are commonly encoded on the DNA strand opposite to the toxin, generally within the coding region or untranslated regions though some are divergently transcribed. The antitoxin binds the toxin mRNA, occluding the binding sites that are necessary for translation ^2–5^ or inducing RNase degradation ^6^. Type I toxins are mostly small, hydrophobic proteins that destabilize cellular membranes at high concentrations, though the exact mechanism of toxicity is not always known ^5,7–10^. Two exceptions are SymE ^3,11^ and RalR ^12^, both nucleases.

One of the first TA systems to be identified was Hok-Sok, a type I system discovered through its ability to stabilize plasmids ^13^ After transcription, the stable Hok mRNA is slowly processed at the 3’ end into a translatable isoform ^14^ This process is attenuated by the highly expressed RNA Sok, which forms a duplex with Hok mRNA leading to subsequent degradation of both transcripts. Should transcription of the TA operon cease, the less stable antitoxin RNA rapidly degrades, allowing unprocessed hok mRNA in the cytoplasm to mature and be translated into a toxic protein that destabilizes cellular membranes ^13,15^.This causes the cell to die or stop replicating upon gene loss, an effect known as post-segregational killing (PSK).

Genes that confer the PSK phenotype, which include some restriction-modification systems, abortive-infection systems, and bacteriocins ^16–18^, are also able to mediate competition between incompatible plasmids. Plasmids segregate to different daughter cells during division and the cells not inheriting the PSK-containing plasmid dies ^19^. Plasmids with PSKs on them are advantaged over non-PSK plasmids ^16,19–21^ accounting for the prevalence of PSK genes on mobile genetic elements (MGEs).

While TA systems were discovered for their effects on plasmids, TAs of all types are also abundant on bacterial chromosomes. The role of TA systems on chromosomes is still uncertain ^22,23^, with theories ranging from them being important components in cellular function to being genomic parasites that persist due to difficulties in displacing them. Proposed cellular functions for the various types of TA systems are mostly stress related, including bacteriostasis, programmed cell death and persister cell formation ^24–26^. Other functions are related to their ability cause PSK:stabilizing genomic regions ^27–29^, neutralizing PSK from plasmid borne TAs ^30^, and acting in abortive infection of bacteriophages ^31,32^ Although some functions have been well characterized for specific loci, such as the ability of type I TA tisB-istR to increase resistance to antibiotics in *E. coli* ^25^, many others have not.

These functions generally rely on the activation of the toxin in response to reductions in transcription or translation of the toxin and antitoxin genes, either because the cell is stressed or because the genes have been lost. This is aided by common characteristics of TA systems, including organization into an operon, antitoxin-mediated regulation of toxin transcription (type II, type III) or translation (type I), and high lability of the antitoxin relative to the toxin ^1 13, 33–35^. Despite shared features, it has been proposed that type I TAs are more likely to be duplicated on chromosomes in a lineage specific manner ^36^ and that they are less mobile than the horizontally promiscuous type II TAs ^17, 30,36–38^. This hypothesis is primarily due to the tendency for type I TAs to found in only a narrow range of species and type II TAs to be more widely distributed.

The mobility of a given gene is affected by certain physical factors, including how it is transferred and post-transfer stablization (e.g. site-specific integration, homologous recombination, replication) ^39^. Yet while any gene may transfer by the above mechanisms, various selective pressures affect which are maintained in populations of descendants. Those that are stably maintained are most likely to be detected during sequencing and subsequent genomic screens. Thus, these screens cannot measure mobility of the genes per se, as genes could be highly mobile but not detected, but can indicated the range of hosts in which the gene has been retained.

We analysed the distribution of type I, type II and type III TA systems across bacterial species and mobile replicons. Consistent with previous claims, type I TAs exhibited a narrow, phyla-specific distribution and were rarer on plasmids than either type II or type III systems. Interestingly, though, this pattern was less consistent on more recently discovered systems. Reasons for these differences, including ability to exhibit PSK, toxin function and gene regulation and biases in the databases and discovery process, are discussed here.

### Type I families occur across a narrow phylogenetic range and are less likely to be on mobile elements than type II and type III families

Within each type of TA system are multiple families of toxins. There is only one corresponding antitoxin for individual type I and III toxins identified so far, but a given type II toxins binds multiple independent families of antitoxins ^38^. We investigated a range of type I, II and III toxin families. Nine type I TA toxin families were included in the analysis (Table 1). All are validated TA systems except for XCV2162 (also known as Plasmid_toxin), which has only been described computationally ^40^. It was included due to its reported distribution, which is consistent with horizontal gene transfer (HGT). SymE is a nuclease, while the remaining are predicted membrane-disrupting proteins. Eleven type II TA toxin families were investigated. Most are part of large, well-described families except for GinA and GinB, which have been described more recently ^38^. Three type III TA families described so far ^41^ were also included.

Phage, plasmid, and bacterial chromosome sequence data were downloaded from the EMBL nucleotide archive (http://www.ebi.ac.uk/ena/, October 2014). These were translated in six frames to derive all possible amino acid sequences from the genomes including the short ORFs that are a characteristic of type I toxins (and traditionally make them difficult to detect). This database was analysed with Profile hidden Markov models (HMMs), derived from the known amino acid sequences for each toxin family as downloaded from Pfam and GenBank. The HMMs for the more recently described families GinA and GinB ^38^ and CptN and TenpN ^41^ were derived from loci reported in the literature.

**Table 1:** Characteristics of type I TA families

| Toxin | Discovery | Phyla:<br>Family | Target | Regulation | References |
| --- | --- | --- | --- | --- | --- |
| Fst | Plasmid<br>stability | 1:6 | Membrane<br>damage | Cis | Weaver *et al.* 2009 |
| Hok | Plasmid<br>stability | 1:4 | Membrane<br>damage | Cis | Gerdes *et al.* 1986 |
| Ibs | Repeats in<br>sequence<br>data | 1:3 | Membrane<br>damage | Cis, | Fozo *et al.* 2008 |
| Ldr | Repeats in<br>sequence<br>data | 1:1 | Membrane<br>damage | Cis | Kawano *et al.* 2002 |
| ShoB | Screening<br>for sRNA | 1:1 | Membrane<br>damage | Trans | Kawano *et al.* 2005 |
| SymE | Screening<br>for sRNA | 5:24 | Ribonuclease | Cis | Kawano *et al.* 2005 |
| TisB | Screening<br>for sRNA | 1:1 | Membrane<br>damage | Trans. | Vogel *et al.* 2004 |
| TxpA | Screening<br>for sRNA | 1:1 | Membrane<br>damage | Cis | Silvaggi *et al.* 2005 |
| XCV2162 | Screening<br>for sRNA | 1:11 | Predicted<br>membrane<br>domain | Cis | Findeiss *et al.* 2010 |

Despite the number of new, unannotated loci found in the family-based searches, type I toxin families were found in fewer phyla than type II and type III families and in fewer species within those phyla (Figure 1). All type I toxins except for SymE, a recently discovered toxin that differs in function from other type I toxins, were found in only one phylum. Some toxins were especially narrow in their distribution:toxins Ldr, ShoB, Txp and TisB were found in less than 5% of species within that phylum (Figure 1) and those species were all in the same family, either Enterobacteriaceae within Proteobacteria or Bacillaceae within Firmicutes (Table 1). None of the most narrowly distributed type I toxin families were found on elements that could be identified as being mobile. The three families found on mobile elements, Fst, Hok, XCV2162, were found in more taxonomic families (six, four, and eleven, respectively) within their phylum (Table 1).

**Figure 1:**
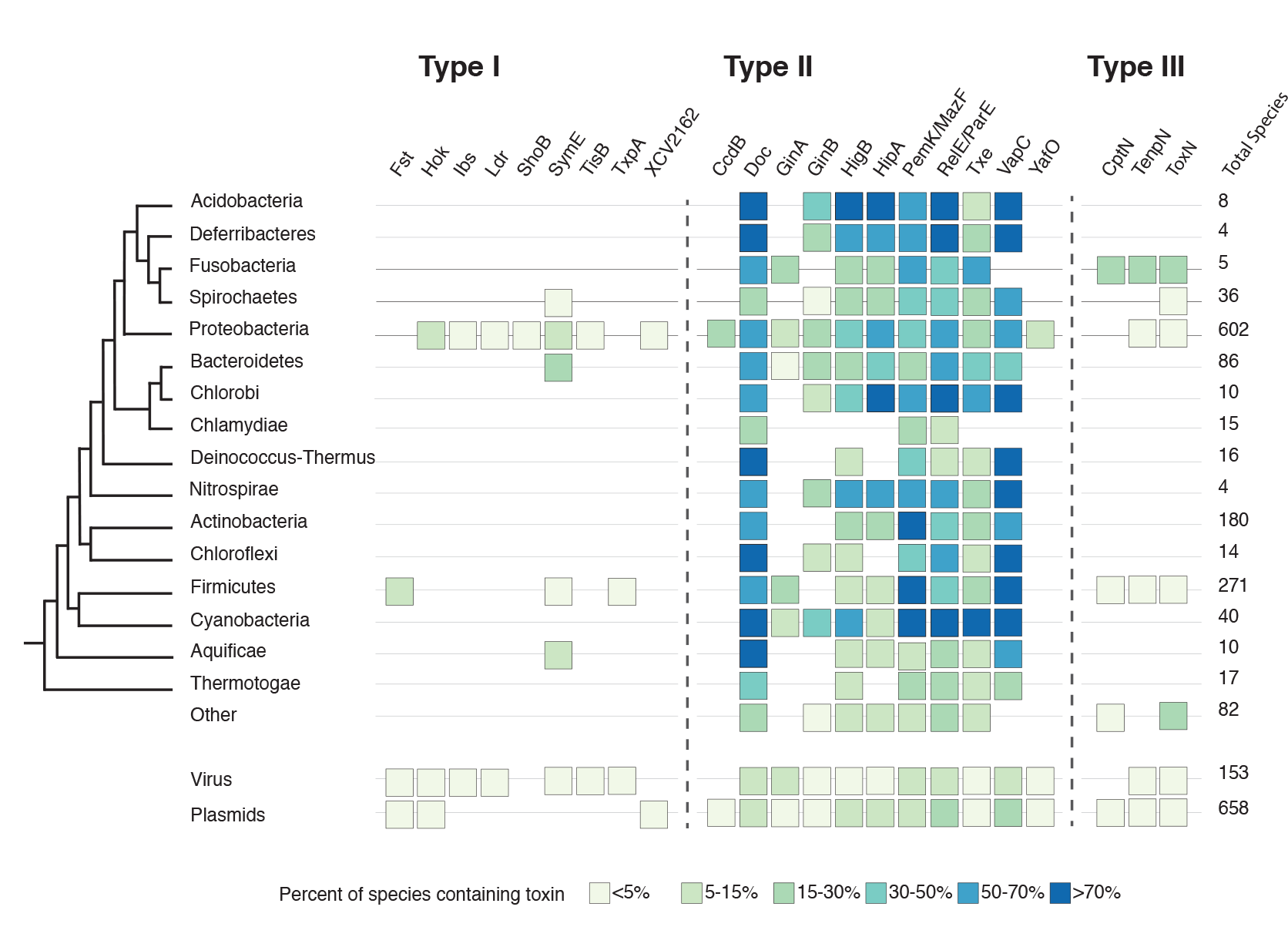
Percent of species within each phyla or replicon that contain a loci from a given type I, type II and type III TA toxin family. HMMs for each TA family were derived from known amino acid sequences and used to search a database of phage, plasmid, and bacterial chromosome sequences subjected to six-frame translations to derive all possible amino acid sequences from that sequence. This includes short ORFs that are typical of type I toxins. For each phyla or replicon, the percent of total species in the database (left of figure) that contain at least one locus of that toxin is reported (boxes).

Most type II families were found across phyla (Figure 1). The families Doc, MazF/PemK, RelE/ParE and VapC were found in all bacterial phyla analyzed, as well as in viruses and plasmids. These TA toxins were prevalent within phyla as well, found on over 70% of species within the phyla Acidobacteria and Cyanobacteria. The two type II toxin families CccdB and YafO were found in only one phylum. Compared to single-phylum type I toxins, these type II toxins were observed in a higher percentage of species, across more taxonomic families within the phylum as well as on mobile elements. Type III families, which also have an RNA antitoxin, were intermediate between type I and type II TAs. All three families were found in both Gram-negative and Gram-positive phyla and on plasmids ^41^, though none were on greater than 30% of the species in the database of translated genomes. As has been reported elsewhere, the distribution of type I toxin families investigated here is comparatively narrow across both species and replicon type. The broader host range of SymE and type III TA systems would suggest the presence of an RNA antitoxin is not the cause. And some type I TAs like Hok and Fst are present on plasmids and capable of HGT, albeit within a narrow host range. Different factors may account for chromosomal-only and narrowly distributed type I families, as discussed below.

### Presence on mobile elements and ability to exhibit PSK as a factor in TA family distribution

In our direct comparison of three types of TA systems, we find that type I TAs are less frequently observed across phyla and on mobile replicons. Generally, we see a correlation between presence on mobile elements and a larger taxonomic range (though the low number of sequenced mobile element prevents showing a strong association) with most type II and III TAs on mobile elements. Except for SymE, those type I TAs found on plasmids are generally in more families than those that are chromosome only (Table 1), though even mobile type I TAs are narrowly distributed compared to type II and III TAs.

Most loci on plasmids, for TAs of all types, are able to mediate PSK ^24, 42^ This is not surprisingly, given the advantage of PSK for the horizontal lifestyle ^19, 20^. However, chromosomal homologues of plasmid-borne TAs often do not exhibit PSK themselves ^30, 43, 44^ A comparison of CcdB family toxins on chromosomes showed that the chromosomally-encoded toxins are under neutral selection, unlike their plasmid-borne homologues ^30^.

Chromosome-only TAs may not be widely distributed on mobile elements simply because they are unable to mediate beneficial phenotypes. Most chromosome-specific type I systems have not been tested for their ability to confer a PSK phenotype (Figure 5), except Ldr-RdlD, which did not confer a PSK phenotype ^31,44^. Because chromosome- and plasmid-borne homologues may be under different selective pressures and have different phenotypes, it is difficult to determine if inability to confer a PSK phenotype is an inherent feature of these chromosome-only TA families, or simply that the particular gene pairs analysed do not show it.

As noted above, carriage on a mobile element does not guarantee distribution across a wide range of host backgrounds. Hok and Fst are on mobile elements, but remain narrowly distributed across phyla. While mobile genes can be selected to have phenotypes that are not beneficial to the host, they still rely on the host for expression of those genes. PSK requires close stoichiometry of toxin and antitoxin, with the toxin remaining inert when the mobile element is in the host and becoming active when the activity of the antitoxin falls below a critical threshold. In cellular backgrounds where the PSK phenotype is neutralized, the genes would no longer be selected ^20^. Families of TA that exhibit PSK in a wide range of cellular backgrounds, may, then, be more likely to be mobilized and more likely to gain entry into new hosts. Expression of the PSK phenotype, or many other phenotypes that may cause the genes to be maintained across a range of hosts, would in turn depend on whether the toxin had a target and whether the genes were appropriately regulated in the new environment.

### Toxin target and regulation of toxins as a factor in TA family distribution

Both toxin target and toxin gene regulation have been proposed as factors in the distribution of TA systems. Type II toxins may be successful across a wide array of species because they affect highly conserved targets ^38^. Most type II toxins are nucleases or gyrase inhibitors ^22, 45^ Type III toxins are all nucleases. Most type I toxins are predicted to be membrane disrupting proteins. Some of these type I systems are toxic in non-related species when expressed at high levels, but may have more specific mechanisms of action when expressed under their native promoter or in a particular genetic background. Lack of target would affect both their ability to express PSK or mediate stress response, and thus could limit their selection across phyla and replicons. It is interesting to note that SymE, a nuclease, is an outlier amongst the type I TAs and is distributed across both Gram negative and Gram positive phyla (though not seen so far on plasmids).

PSK, abortive infection and TA-mediated stress response all require that the toxin is inactive but can be released upon specific stimuli. Expression of the toxin and its cognate antitoxin must be tightly regulated within the cell. Small changes in gene regulation were believed to be why *Bacillus subtilis* could be used to amplify clones of only some of the Fst loci from various bacterial species:others appeared to cause cell death when moved into the new cellular background ^33^.

While type I TAs do differ in their regulation from type II and type III, how this may affect their distribution is still a matter for speculation. Regulation of type I TAs occurs at the RNA level:free mRNA must be translated to produce a toxin ^46^. In type II and III systems the antitoxin (protein or RNA) directly binds the already-translated toxin these systems and pools of toxin remain in the cytoplasm at all times. This may make type III systems, for example, more suited to responding to phage infection, where the toxin can be quickly released ^46^. Within type I TAs, most toxins are regulated by an RNA encoded on the opposite strand to the toxin ^5,29^. All plasmid-borne type I antitoxins follow this pattern. Two toxins described here, TisB and ShoB, have RNA antitoxins transcribed from a different locus (as do other toxins Zor and DinQ ^47,48^. They are particularly narrowly distributed, found in only one taxonomic family. These RNAs often have smaller regions of complementarity and some required additional components to stabilize the interactions (although Hfq, which fulfils this function, is not known to be necessary for any type I TA regulation other than RalR ^29^). As more type I TAs are discovered, stronger patterns of regulation and how this relates to host range may become apparent.

### Biases in databases and discovery of type I TA families may account for apparent narrowness of type I distribution

It can be difficult to make sweeping statements on the distribution of any gene across many phyla and mobile elements due to selection of sequenced genomes and how well bioinformatics tools find different genetic elements within them. Proteobacteria and Firmicutes are the two most studied phyla of bacteria with the greatest number of sequenced genomes, and contain a disproportionate number of TA systems (Figure 1 and S1). This is particularly true when strain rather than species is analyzed, or when the number of species within a phylum that has the toxin is not normalized by total number of species within that phyla (Figure S1).

Type I TA systems have historically been more difficult to detect in silica, with sequence-diverse RNA antitoxins and small toxins (under 60 amino acids), potentially reducing both the number of toxin families we know of and the number of phyla in which they have been found. Many type II families reported here are actually super-families of many described toxins. Aggregating these can increase their apparent distribution. CcdB has a narrow distribution in this screen, but is often combined with MazF and related families, making the super-family relatively widely distributed. Type I toxins Ldr and Fst are considered to be related ^36,38^ and combining them together would result in a broad family. The apparent narrowness of type I TA systems, then, may be a result of bioinformatic limitations.

The use of sequences and other information derived from known TA systems to screen for new TA systems also imposes an obvious bias onto the search. Many of the best-studied and most widely distributed type II systems were discovered due to a phenotype, usually the ability to stabilise plasmids in monocultures. It is not surprising, then, that these families were later found to be widely distributed on mobile elements. On the other hand, many of the type I systems discovered were first described on chromosomes. Ultimately, methods that go beyond sequence-based features of known TA systems are more likely to yield families with novel characteristics. The narrowly distributed type II families YafO, GinC and GinD were discovered bioinformatically ^38^ due to their association with known antitoxins (guilt by association) rather than sequence features of the toxin. They exhibited patterns of distribution similar to many type I families (Figure 1). Another approach which may yield novel types of TA systems is that of Sberro *et al.* ^49^. This group studied genomes that had been randomly fragmented and inserted into *E. coli* plasmids for sequencing. They identified genes that were only present on fragments (and thus could be amplified in *E. coli*) when an adjacent ORF was present, implying a toxin and antitoxin function. These were filtered for genes that appeared as homologues across species, suggesting HGT, and in regions of the genome associated with phage defence to find novel TA systems ^49^.

## Conclusion

We found that type I toxin families are less often found on known mobile elements or distributed across large taxonomic ranges when compared to type II and type III families. It has frequently been suggested that type I TAs are more lineage specific than type II TA systems. Howeverer, the broader phylogenetic distribution of type I toxin SymE and type III TA systems seen here would suggest one of the defining features of type I systems, the presence of an RNA antitoxin, does not account for the difference.

The factors that select for maintenance of horizontally acquired genes vary. Genes on both chromosomes and plasmids can be selected by within-host forces ^21^. All TA systems consist of a tightly regulated antitoxin and toxin capable of stopping bacterial growth-some systems even have reversible effects, able to be turned off when conditions change. These make for versatile modules with the potential to fill a variety of functions, from plasmid competition (eg, PSK) to cellular stress response to phage-plasmid competition (eg abortive infection). The function of a given locus will depend in part on its history, though some families may have features that make them more able to fill certain roles. It could be that aspects of the toxin target and gene regulation affect the ability of some type I TAs to stably establish in new species. Many narrowly distributed type I TAs are membrane proteins, and the most narrowly distributed (along with one narrowly distributed type II TA) are integrated into host stress networks. The only type I nuclease studied here was also the only type I toxin to be found across phyla. All type III and many type II toxins are nucleases.

Alternatively, as our ability to detect and analyze type I TAs increases, these patterns may become weaker. The families found so far may remain chromosomal and/or narrowly distributed, despite an increase in genome and mobile element sequencing. But new types and new families of TAs within those types are being described at a rapid pace ^37, 46, 50, 51^, some of which have alternative distributions. We see this already, with type I TA toxins SymE and RalR, which are nucleases and type II toxins like YafO, GinB and GinC that are narrowly distributed. The potential type I TA XCV2162 is plasmid-borne and has an erratic phylogenetic distribution consistent with HGT ^40^. It may be that the patterns of lineage dependence so far attributed to type I TAs as a group will turn out to be a feature of specific families within all types of TAs, and it is only that we found the narrowly-distributed, membrane associated families of type I TAs and the broadly distributed central-dogma targeting type II TAs first.

## Acknowledgements

The authors would also like to thank the Marsden Fund M1138, the McCarthy Fund and the NIH for travel funds during the extent of this research.

## Supplementary Information

**Figure S1:**
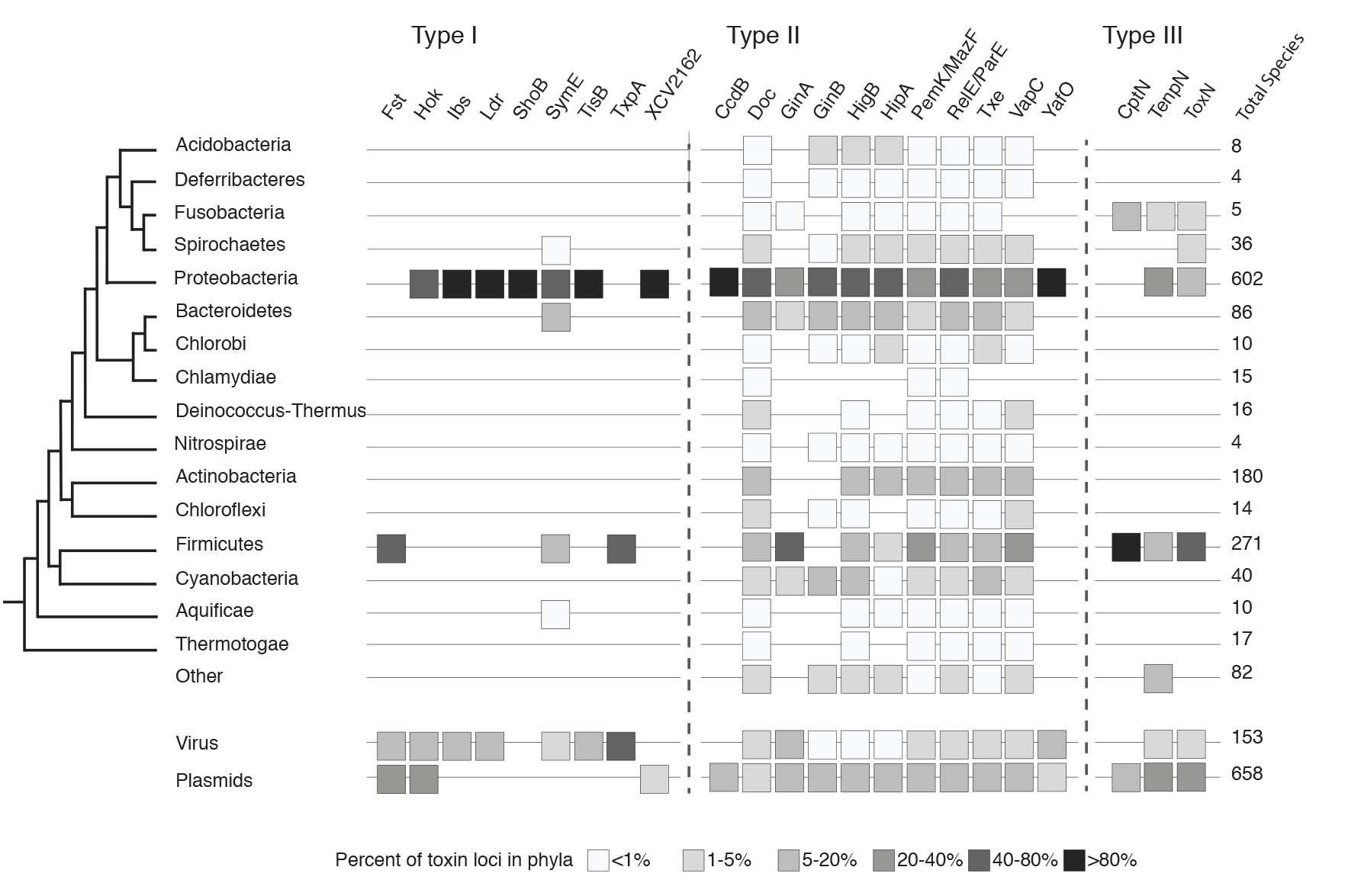
Percent of total toxin loci for type I, type II and type III TA toxin families found in a given phyla or replicon. HMMs for each TA family were derived from known amino acid sequences and used to search a database of phage, plasmid, and bacterial chromosome sequences subjected to six-frame translations to derive all possible amino acid sequences from that sequence. This includes short ORFs that are typical of type I toxins. The percent of all species bearing that toxin that were found in a given phyla is reported (boxes).

**Table S1:**
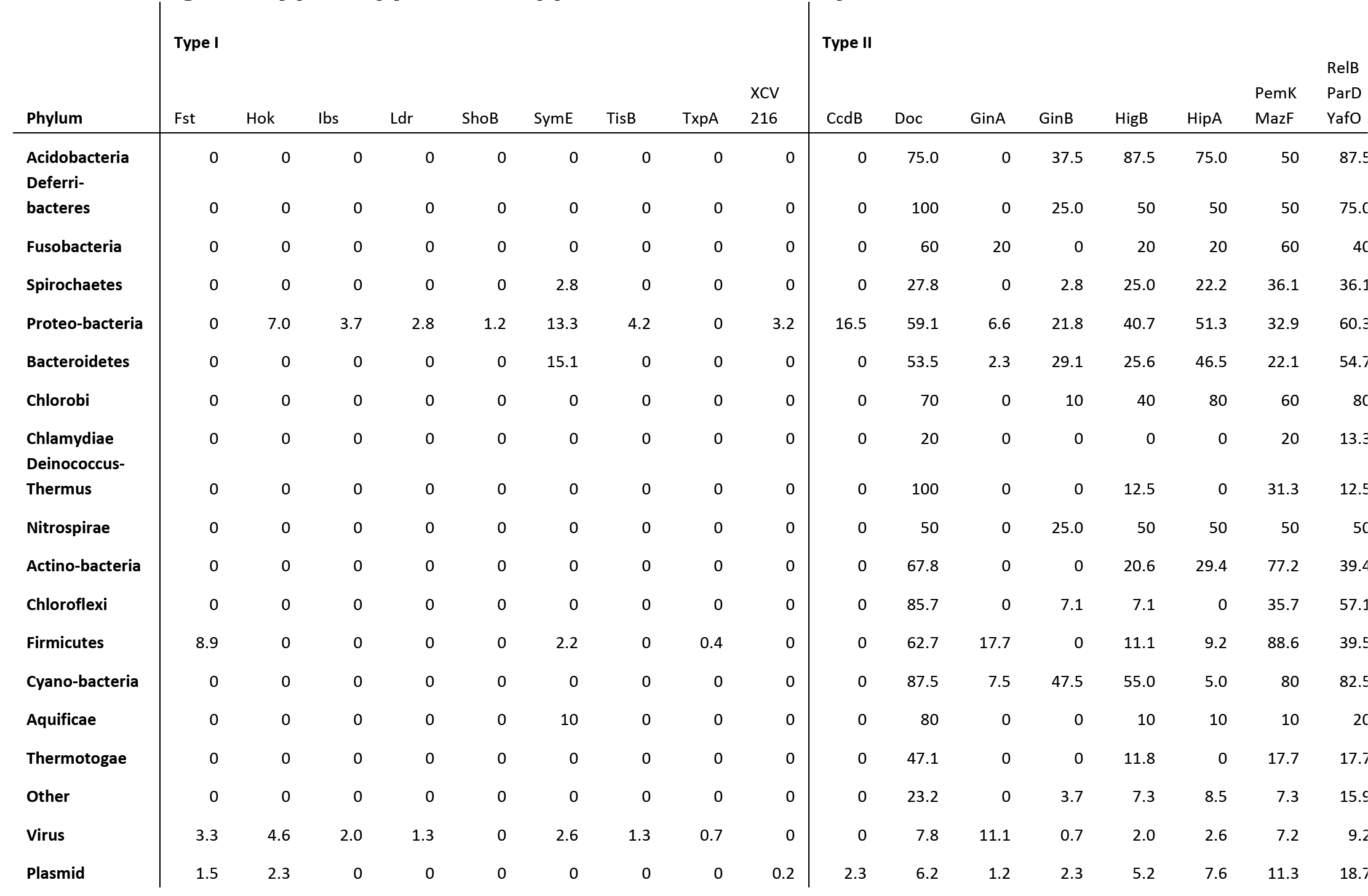
Percent of species within each phyla or replicon that contain aloci from a given type I, type II and type III TA toxin family

**Table S2:**
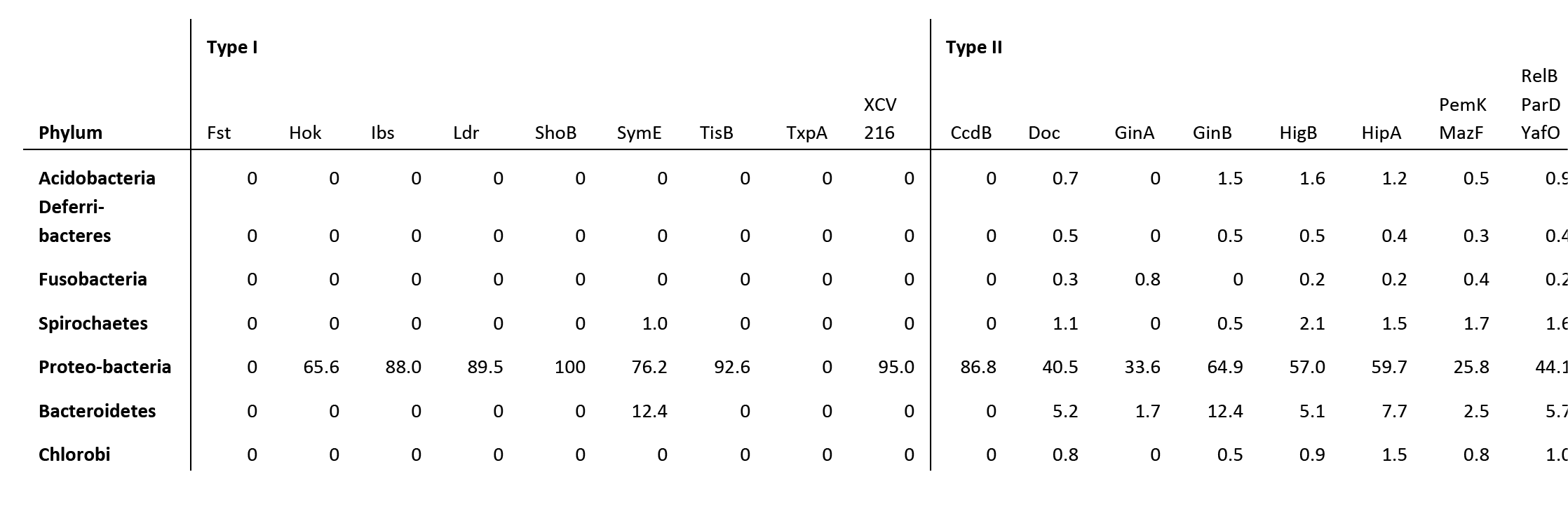
Percent of total toxin loci for type I,stype II and type III TA toxin families found in a given phyla or replicon

|  | Type I |  |  |  |  |  |  |  |  |  | Type II |  |  |  |  |  |  |
| --- | --- | --- | --- | --- | --- | --- | --- | --- | --- | --- | --- | --- | --- | --- | --- | --- | --- |
| Phylum | Fst | Hok | Ibs | Ldr | ShoB | SymE | TisB | TxpA | XCV<br>216 | CcdB | Doc | GinA | GinB | HigB | HipA | PemK<br>MazF | RelB<br>ParD<br>YafO |
| Acidobacteria | 0 | 0 | 0 | 0 | 0 | 0 | 0 | 0 | 0 | 0 | 0.7 | 0 | 1.5 | 1.6 | 1.2 | 0.5 | 0.9 |
| Deferri-<br>bacteres | 0 | 0 | 0 | 0 | 0 | 0 | 0 | 0 | 0 | 0 | 0.5 | 0 | 0.5 | 0.5 | 0.4 | 0.3 | 0.4 |
| Fusobacteria | 0 | 0 | 0 | 0 | 0 | 0 | 0 | 0 | 0 | 0 | 0.3 | 0.8 | 0 | 0.2 | 0.2 | 0.4 | 0.2 |
| Spirochaetes | 0 | 0 | 0 | 0 | 0 | 1.0 | 0 | 0 | 0 | 0 | 1.1 | 0 | 0.5 | 2.1 | 1.5 | 1.7 | 1.6 |
| Proteo-bacteria | 0 | 65.6 | 88.0 | 89.5 | 100 | 76.2 | 92.6 | 0 | 95.0 | 86.8 | 40.5 | 33.6 | 64.9 | 57.0 | 59.7 | 25.8 | 44.1 |
| Bacteroidetes | 0 | 0 | 0 | 0 | 0 | 12.4 | 0 | 0 | 0 | 0 | 5.2 | 1.7 | 12.4 | 5.1 | 7.7 | 2.5 | 5.7 |
| Chlorobi | 0 | 0 | 0 | 0 | 0 | 0 | 0 | 0 | 0 | 0 | 0.8 | 0 | 0.5 | 0.9 | 1.5 | 0.8 | 1.0 |

|  |  |  |  |  |  |  |  |  |  |  |  |  |  |  |  |  |  |
| --- | --- | --- | --- | --- | --- | --- | --- | --- | --- | --- | --- | --- | --- | --- | --- | --- | --- |
| **Chlamydiae** | 0 | 0 | 0 | 0 | 0 | 0 | 0 | 0 | 0 | 0 | 0.3 | 0 | 0 | 0 | 0 | 0.4 | 0.2 |
| **Deinococcus-**<br>**Thermus** | 0 | 0 | 0 | 0 | 0 | 0 | 0 | 0 | 0 | 0 | 1.8 | 0 | 0 | 0.5 | 0 | 0.7 | 0.2 |
| **Nitrospirae** | 0 | 0 | 0 | 0 | 0 | 0 | 0 | 0 | 0 | 0 | 0.2 | 0 | 0.5 | 0.5 | 0.4 | 0.3 | 0.2 |
| **Actino-bacteria** | 0 | 0 | 0 | 0 | 0 | 0 | 0 | 0 | 0 | 0 | 13.9 | 0 | 0 | 8.6 | 10.2 | 18.1 | 8.6 |
| **Chloroflexi** | 0 | 0 | 0 | 0 | 0 | 0 | 0 | 0 | 0 | 0 | 1.4 | 0 | 0.5 | 0.2 | 0 | 0.7 | 1.0 |
| **Firmicutes** | 61.5 | 0 | 0 | 0 | 0 | 5.7 | 0 | 50 | 0 | 0 | 19.3 | 40.3 | 0 | 7.0 | 4.8 | 31.3 | 13.0 |
| **Cyano-bacteria** | 0 | 0 | 0 | 0 | 0 | 0 | 0 | 0 | 0 | 0 | 4.0 | 2.5 | 9.4 | 5.1 | 0.4 | 4.2 | 4.0 |
| **Aquificae** | 0 | 0 | 0 | 0 | 0 | 1.0 | 0 | 0 | 0 | 0 | 0.9 | 0 | 0 | 0.2 | 0.2 | 0.1 | 0.2 |
| **Thermotogae** | 0 | 0 | 0 | 0 | 0 | 0 | 0 | 0 | 0 | 0 | 0.9 | 0 | 0 | 0.5 | 0 | 0.4 | 0.4 |
| **Other** | 0 | 0 | 0 | 0 | 0 | 0 | 0 | 0 | 0 | 0 | 2.2 | 0 | 1.5 | 1.4 | 1.4 | 0.8 | 1.6 |
| **Virus** | 12.8 | 10.9 | 12.0 | 10.5 | 0 | 3.8 | 7.4 | 50 | 0 | 0 | 1.4 | 14.3 | 0.5 | 0.7 | 0.8 | 1.4 | 1.7 |
| **Plasmid** | 25.6 | 23.4 | 0 | 0 | 0 | 0 | 0 | 0 | 5.0 | 13.2 | 4.7 | 6.7 | 7.4 | 7.9 | 9.7 | 9.7 | 14.9 |

